# The flow state is not accompanied by frontal-midline theta activity: An EEG investigation of more than 700 video gameplay sessions

**DOI:** 10.1101/2024.07.11.603158

**Authors:** Hirotaka Sugino, Takuya Ideriha, Ryoichiro Yamazaki, Junichi Ushiyama

## Abstract

People sometimes experience a “flow state”—characterized by hyperfocus, time distortion, and loss of self-awareness—during sports or video gameplay. Previous neuropsychological studies using simple laboratory tasks have reported that the flow state is associated with activation in the frontal lobe, reflected in theta (4–7 Hz) band rhythmic neural activity in medial prefrontal regions (frontal-midline theta [FMT] activity). However, the findings of previous studies might be problematic because they did not appropriately capture the neural activity associated with the flow state for the following reasons: 1) they used unfamiliar and unmotivating tasks; 2) they defined the neural basis of the flow state as neural activity occurring during tasks of optimal difficulty, disregarding trial-to-trial variations in subjective experience of the flow state; 3) the duration of the experiment or the number of trials was not sufficient to capture the rare experience of flow; or 4) they ignored individual differences in neural activities related to flow experiences. Thus, we examined the relationship between the flow state and FMT activity, recorded via scalp electroencephalography, in an experimental paradigm that addressed these four issues. First, participants played their favorite competitive video games, which they had been routinely playing. Second, task difficulty was kept as uniform as possible across trials by employing rank matching to directly examine the correlation between subjective flow level and FMT activity across trials. Third, to address the concern regarding the low frequency of the flow experience, more than 100 trials were completed over 10 days by each participant. Lastly, we adopted a within-participant statistical approach to examine individual differences in the nature of the flow experience. The results showed no correlation between FMT activity and the degree of subjective flow in six out of seven participants, contrary to previous reports. Our results challenge the conventional view that frontal lobe activity, as reflected in FMT activity, is instrumental in entering into the flow state.

## 1. Introduction

People sometimes experience a distortion of time or loss of self-awareness when intensely focusing on a task. This phenomenon is referred to as the “flow state,” which is defined as “a psychological state in which the person feels simultaneously cognitively efficient, motivated, and happy” (Moneta and Csikszentmihalyi 1999). It has been reported that the subjective experience of flow is accompanied by a number of unique affective and attentional states, such as strong motivation, high energy, immersion, and complete attentional focus (Moneta and Csikszentmihalyi 1999; Gold and Ciorciari 2020). Additionally, the association between improved performance and the flow state has also been observed in sports (Stavrou et al. 2007), music (MacDonald, Byrne, and Carlton 2006), and video gameplay (Jin 2012).

The neural mechanisms of the flow state are currently under study. Some electroencephalography (EEG) studies have proposed that theta (4–7 Hz) band oscillation in the medial prefrontal region is the neural correlate of the flow state (Ewing, Fairclough, and Gilleade 2016; Katahira et al. 2018; Knierim et al. 2018; Farrugia et al. 2021). Such rhythmic neural activity, known as frontal-midline theta (FMT) activity, is of significant interest as a potential neural marker for top-down cognitive control and attention (Cavanagh and Frank 2014; Berger et al. 2019; Pavlov and Kotchoubey 2020). It is assumed that intense recruitment of top-down cognitive networks, as indexed by strong FMT activity, leads to hyperfocus and low self-awareness during flow (van der Linden, Tops, and Bakker 2021). For instance, Ewing et al. (Ewing, Fairclough, and Gilleade 2016) observed higher FMT activity during video game playing under optimal difficulty conditions versus conditions in which the game was rated either too easy or too hard; this was confirmed by questionnaires.

Several other studies also have attempted to reveal the neural mechanisms of the flow state using such an experimental design, comparing neural activities between tasks of optimal difficulty and tasks with other difficulty levels (Ewing, Fairclough, and Gilleade 2016; Katahira et al. 2018; Knierim et al. 2018; Farrugia et al. 2021; Ulrich et al. 2014; Yoshida et al. 2014). However, Alameda et al. (Alameda, Sanabria, and Ciria 2022) pointed out limitations of the experimental design employed in studies attempting to elucidate the neural correlates of the flow state by comparing neural activity during optimally difficult tasks with that during easier or more difficult tasks. Their article criticized these studies for ignoring the fluctuation in the flow state that may occur even under a constant level of difficulty. In fact, even in sports involving the repetition of a single movement, such as swimming or sprinting, the difficulty of which theoretically does not fluctuate, it is not always possible for athletes to enter the flow state in competitions. Because of the inherent uncertainty in the experience of flow, it is natural to assume that the degree of flow varies on a trial-by-trial basis within an individual even when task difficulty is constant. Thus, to prevent the fluctuation in flow state between tasks that are identical in difficulty from being ignored, paradigms interpreting neural activity during optimally difficult tasks as the neural correlate of the flow state should not be used. Additionally, because cognitive load (e.g., task difficulty) is known to influence neural activities, including FMT activity (Onton, Delorme, and Makeig 2005; Popov et al. 2018), the neural correlates reported by previous studies might be related not only to the flow state but also to task difficulty.

Moreover, the infrequency of flow experiences was also noted as a prominent issue in previous studies (Alameda, Sanabria, and Ciria 2022). Even elite athletes, whose capacity to experience the flow state is considered greater because of the familiarity of the tasks they undertake and their strong motivation, do not always experience flow. However, previous studies investigating the flow state often adopted experimental procedures in which participants were engaged in unfamiliar and unmotivating tasks for only a few dozen minutes. Given the infrequency of flow experiences, such procedures would be unlikely to induce flow during tasks.

In addition to the issues pointed out by Alameda et al. (Alameda, Sanabria, and Ciria 2022), the statistical approach in previous studies was not ideal to capture individual differences in the nature of the flow state. In other words, such studies examined the neural basis of the flow state at the group level, ignoring individual differences. Psychological and emotional states vary considerably among individuals. Therefore, methodologies that explored the neural mechanisms of psychological states on the basis of group averages might have drawn erroneous conclusions (Fischer, Nilsson, and Ebner 2023; Smith and Little 2018). Recently, problematic conclusions of studies calculating the group averages of psychological indices have been reported (Grice et al. 2020). For example, although Siegel et al. (Siegel et al. 2018) obtained an adequate effect size to support the hypothesis of their psychological experiment, a reanalysis of their data by Grice et al. (2020) revealed that only 11 of 45 participants showed results consistent with the hypothesis (Grice et al. 2020). Therefore, to avoid misleading conclusions, it has been suggested that a within-participant statistical approach that also evaluates the proportion of the population that show a given outcome is a preferable alternative approach (Grice et al. 2020).

Here, we investigated whether FMT activity correlates with subjective flow states in an experiment designed to resolve the issues in previous studies described previously. The unique features of this study are as follows: 1) participants played their favorite video games (in which they were also most proficient) to facilitate the experience of the flow state; 2) participants consistently played the games at an appropriate difficulty level (e.g., by engaging in rated matches) throughout the experiment; 3) each participant completed more than 100 gameplay sessions, with EEG recording, within 10 days to overcome the low frequency of flow experiences; and 4) we adopted a within-participant statistical approach to capture individual differences in flow states, and the correlation between subjective flow level and FMT activity was calculated across trials.

## 2. Methods

### 2.1. Participants

Seven healthy individuals (six males and one female with a mean [standard deviation] age of 24 [2.89] years on the first day of the experiment) participated. None of the participants had experienced any neurological disorders. Informed consent was obtained from all participants prior to the experiment. All research procedures were approved by the Research Ethics Committee in Shonan Fujisawa Campus, Keio University (approval number: 334) and were conducted in accordance with the Declaration of Helsinki, except that the study was not pre-registered in a database.

### 2.2. Recording

#### 2.2.1. Electroencephalogram recordings

Scalp EEG was recorded from 19 channels (Fp1, Fp2, F7, F3, Fz, F4, F8, T3, C3, Cz, C4, T4, T5, P3, Pz, P4, T6, O1, O2) according to the international 10–20 system of electrode placement. Passive Ag/AgCl electrodes 18 mm in diameter were mounted in an elastic cap (g.GAMMAcap 1027; Guger Technologies, Graz, Austria). The reference and ground electrodes were placed on the left and right earlobe, respectively. EEG signals were amplified and bandpass filtered at 0.5–200 Hz using a biosignal recording system (g.BSamp 0201a; Guger Technologies). A 50-Hz digital notch filter was applied to eliminate artifacts stemming from alternating current line noise. All analog EEG signals were digitized at a sampling rate of 1,000 Hz through a 16-bit resolution analog-to-digital converter (NI 779987-01; National Instruments, Austin, TX, USA), controlled by data-logging software that was custom built using MATLAB software (version 9.14.0.2206163 [2023a]; MathWorks, Inc., Natick, MA, USA) running on a laptop PC (G-Tune P5; Mouse Computer Co., Ltd., Tokyo, Japan).

#### 2.2.2. Electrocardiogram recordings

To record electrocardiograms (ECGs), three disposable 24-mm-diameter Ag/AgCl electrodes (Kendall H124SG; Cardinal Health, Inc., Dublin, OH, USA) were placed on the upper segment of the sternum, on the ensiform process, and below the left clavicle (negative, positive, and ground electrodes, respectively) (NASA lead method; (Takaya et al. 2021). The acquired ECG signals were amplified and bandpass filtered at 5–500 Hz with the same amplifier used for the EEG recording system mentioned above, resulting in both signals being synchronized in time without line noise.

#### 2.2.3. Psychological questionnaires

Participants completed the Flow Short Scale (FSS; Engeser and Rheinberg 2008), a questionnaire evaluating the subjective flow experience and the balance between challenge and skill (Engeser and Rheinberg 2008). The FSS consists of 10 items pertaining to flow experiences and 3 pertaining to the challenge–skill balance. Seven- and nine-point Likert scales, ranging from 1 to 7 and 9, are used to measure the subjective flow experience and balance, respectively (see Appendix). Participants completed all 13 items immediately after game playing in each trial.

### 2.3. Procedures

#### 2.3.1. Experimental design

Each participant came to the laboratory on 10 non-consecutive days and completed the experiment on an individual basis. On a single day, the experimental procedure comprised baseline and task phases. In the baseline phase, resting state EEG and ECG were recorded for 2 minutes with participants’ eyes open. Following the baseline phase, participants completed at least 10 trials in the task phase. Each trial started with the playing of a video game (see “Details of Game Playing” section for comprehensive information), during which EEG and ECG were recorded using the same procedure as in the baseline phase. After game playing, participants completed psychological questionnaires. The task phase trials continued until (1) the participant met the criterion for success in three consecutive trials (from the eighth trial onward) or (2) 90 minutes had passed since the first trial. The former condition aimed to motivate participants to perform well until the end of the task phase. The latter condition was established to avoid participant fatigue. Each experiment was < 2 hours in duration on any given day. Throughout the experiment, participants sat on a comfortable chair. To prevent artifacts contaminating the data, the participants were instructed to avoid muscle contractions unnecessary for playing a game.

#### 2.3.2. Details of game playing

Each participant played the game that was both his/her favorite and the one he/she was most confident about in terms of his/her skill. Table 1 summarizes the games and hardware selected by each participant. The methodology for controlling the difficulty of trials in the task phase differed among game genres. For fighting games (three participants), shooter games (two participants), and racing games (one participant), participants were automatically matched with online player(s) with an equivalent rank. For a rhythm game, chosen by a single participant, the participant selected and played songs with a level of difficulty that allowed her to match all notes approximately once in every two attempts. The criteria for success in a trial during the task phase were defined according to game genre. For fighting and shooter games, a successful trial was defined as victory in a match. For a racing game, finishing a race in sixth place or higher (field of 12 racers) was judged as successful. For the rhythm game, matching all notes in a song was defined as success. These criteria were designed to keep the success rate at approximately 50%, which is considered to represent a good balance between difficulty and participant skill. It is important to note that all participants played the games via their own user accounts such that poor performance (e.g., losing or playing no role in a match) led to losses that lowered their actual online rank. This prevented participants from intentionally making mistakes or playing carelessly.

**Table 1.** Video games used in the experiment.

| Participant ID | Game title | Game type | Hardware |
| --- | --- | --- | --- |
| 1,2,3 | Super Smash Bros. Ultimate | Fighting | Nintendo Switch® (Nintendo Co., Ltd., Kyoto, Japan) |
| 4 | Mario Kart 8 Deluxe | Racing | Nintendo Switch® (Nintendo Co., Ltd., Kyoto, Japan) |
| 5 | Splatoon 3 | Third-person shooter | Nintendo Switch® (Nintendo Co., Ltd., Kyoto, Japan) |
| 6 | VALORANT | First-person shooter | Custom-built PC (Windows 11, 13th Intel® Core™ i7-13700F; Project White Co., Ltd. Gumma, Japan) |
| 7 | BanG Dream! Girls Band Party! | Rhythm game | iPhone SE (2022, Apple, CA, USA) |

### 2.4. Data processing and analysis

#### 2.4.1. EEG processing

The recorded EEG signals were processed and analyzed using MATLAB scripts and the EEGLAB toolbox (version 2023.0)(Delorme and Makeig 2004). The EEG signals were low-pass filtered at 100 Hz, high-pass filtered at 3 Hz, and band-stop filtered at 49–51 and 99– 101 Hz using a third-order Butterworth filter. These filtered signals were segmented according to an epoch length of 1,000 ms. After removing epochs containing apparent artifacts (detected by visual inspection), the remaining data were decomposed with independent component analysis to remove components related to eye blinks and body movements. The decomposed data were used for computing power spectrum density (PSD). Welch’s method was used to calculate the PSD by applying a Hanning window of 1,000 ms with zero overlap. The ratio of theta PSD (4–7 Hz) to PSD of frequencies < 50 Hz (i.e., 1-49 Hz), called the theta ratio, was calculated for each electrode and trial. The theta ratio of the Fz electrode was adopted as an index of FMT activity, defined as the FMT ratio.

#### 2.4.2. ECG processing

MATLAB scripts automatically detected R-peaks in the obtained ECG signals. Visual inspection confirmed the peaks. When this inspection process revealed errors in R-peak detection, the experimenter manually corrected them. The temporal interval between two adjacent R-peaks, the R–R interval (RRI), was calculated for all R-peaks. Even though many previous studies dealing with heart rates averaged RRI values for each trial to evaluate sympathetic activity (Järvelin-Pasanen, Sinikallio, and Tarvainen 2018), such mean RRI values may be affected by dynamic trends in the data. Therefore, this study did not use mean RRI values, instead adopting the ratio of the minimal RRI in each trial to that of the corresponding baseline (RRI_min_) as an index of sympathetic activity (Vicente-Rodríguez et al. 2020). Note that the purpose of this calculation was to eliminate the effect of circadian rhythms (Catai et al. 2020).

#### 2.4.3. Processing of psychological and behavioral data

The scores for the 10 FSS items assessing subjective flow experience were averaged for each trial as an index of the depth of the subjective flow state (“flow index”). Meanwhile, the score for the item pertaining to the subjective difficulty of each trial was defined as the “difficulty index.” The game playing performance on each trial was classified as success or failure, as determined by the previously described criteria.

### 2.5. Statistical Analysis

We collected 1009 trials in total. We excluded 9 trials from analyses due to equipment malfunction in data recording. For the same reason, we excluded baseline recordings for 5 days.

Accordingly, we excluded 73 trials for such days without baseline data. Also, we rejected 5 trials in which participants did not answer the FSS questionnaires correctly. Eventually, 920 trials remained for the analyses.

The Mann–Whitney U test was used to compare the FMT ratio for all baseline recordings with that for all trials. Because the linearity of the flow index was not guaranteed, Spearman correlation of the flow index was performed using the RRI_min_ and the FMT ratio. In addition, trials were divided into three bins according to the difficulty index (“optimal bin” for trials scoring 5; “easy bin” for trials scoring < 5; “hard bin” for trials scoring > 5) and into two bins according to the behavioral performance (“success bin” and “failure bin”). The Mann– Whitney U test was used to examine differences according to the difficulty and performance bins. For multiple comparison correction, the false discovery rate was controlled by applying the Benjamini–Hochberg method (Benjamini and Hochberg 1995) to the *p*-values of all statistical tests for each participant.

All statistical analyses were applied using a within-individual approach (Grice et al. 2020). The present study focused on the number of individuals showing statistically significant results and the wider trends in the results.

## 3. Results

### 3.1. EEG: FMT activity during flow states

#### 3.1.1. Comparison with the resting state

Table 2 summarizes the descriptive statistics of indices in the task phase (i.e., the FMT ratio, RRI_min_, and flow and difficulty indices) for each participant. Five out of the seven participants showed a significantly higher FMT ratio during games than at rest (Fig. 1, Table 3). Note that the remaining two participants played racing and first-person shooter games, confirming the absence of an association between the observed increase in the FMT ratio and game genre. These results suggest individual differences in the neural basis of the flow state during video game playing, i.e., the top-down network centered in the frontal lobe is required for flow while playing games in most, but not all people.

**Figure 1.**
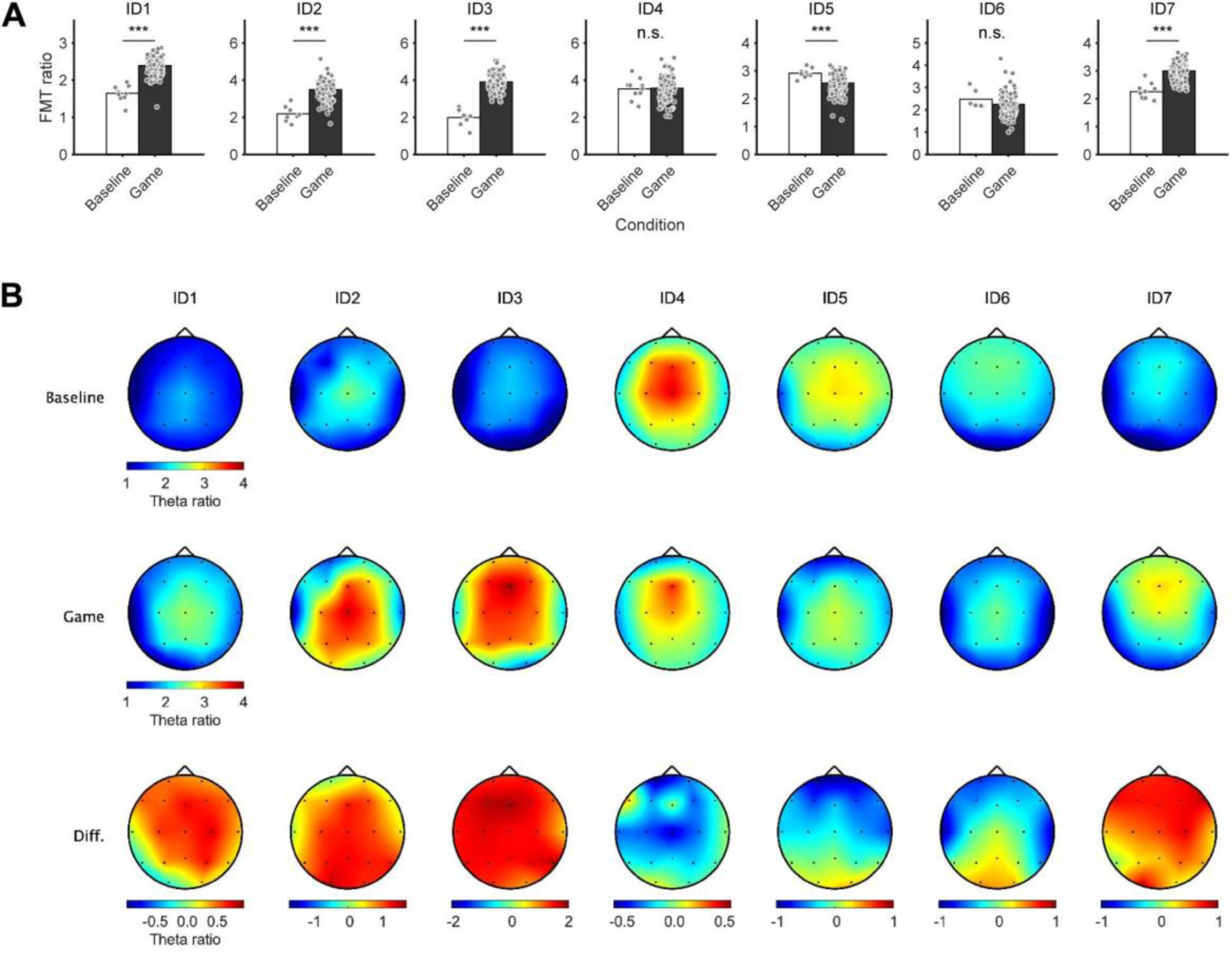
Comparison of frontal-midline theta (FMT) activity between the gameplay and baseline phases within individuals. **(A)** Bars represent the FMT ratio between the gameplay (black) and baseline (white) phases. Each dot denotes one trial. The false discovery rate was corrected for multiple comparisons. ***p < 0.001; n.s., not significant. **(B)** Topography of theta power in the baseline (top) and gameplay (middle) phases, and the difference in power between phases (game - baseline; bottom). Each column corresponds to one participant.

**Table 2.**
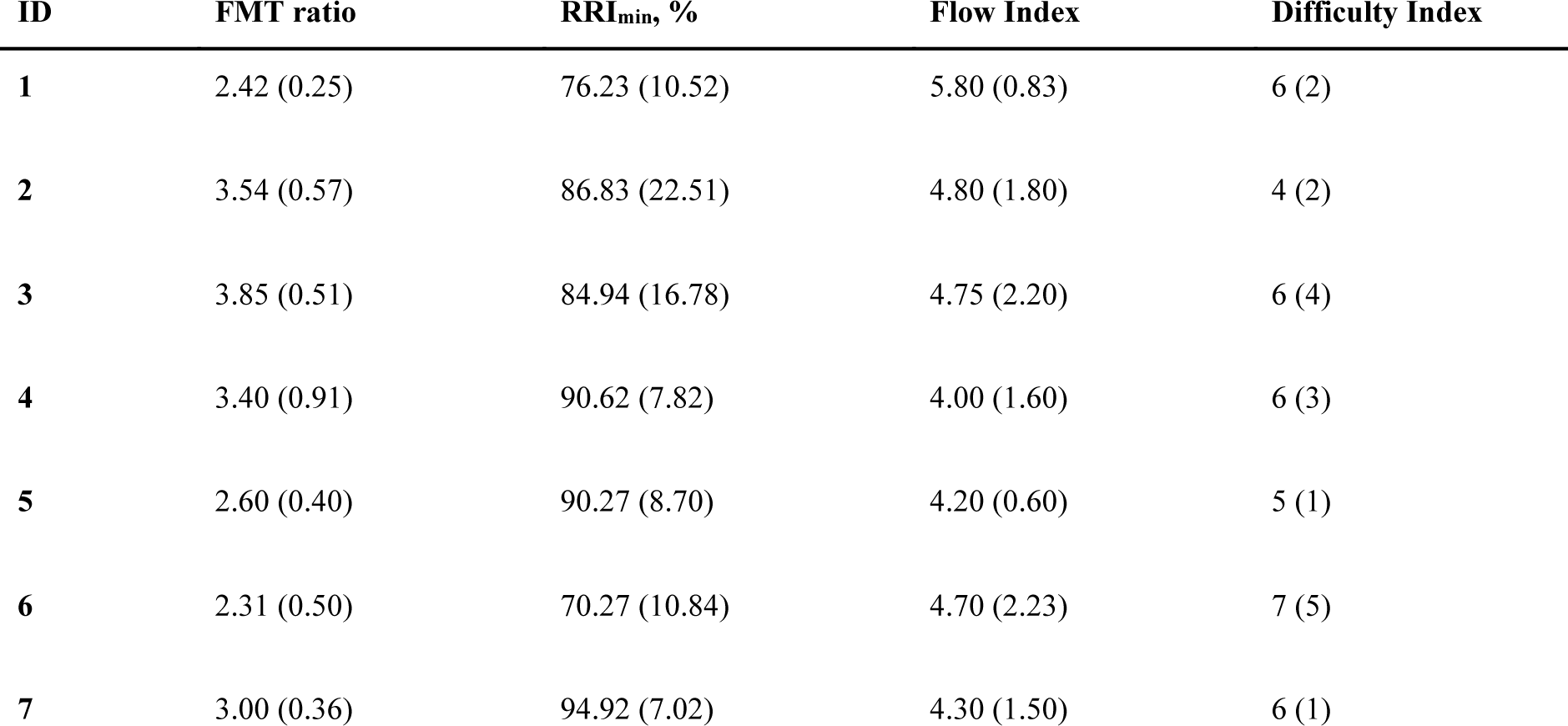
Descriptive statistics of indices in the task phase for each participant. The values are medians (interquartile range).

**Table 3.** Results of statistical analyses of the frontal-midline theta (FMT) ratio. The false discovery rate was corrected for multiple comparisons. *p < 0.05; **p < 0.01; ***p < 0.001.

| FMT ratio |  |  | ID1 | ID2 | ID3 | ID4 | ID5 | ID6 | ID7 |
| --- | --- | --- | --- | --- | --- | --- | --- | --- | --- |
| Gaming vs. Baseline |  | Z | -5.21 | -4.48 | -4.75 | -0.31 | 3.30 | 0.95 | -4.90 |
|  |  | p | < 0.001*** | < 0.001*** | < 0.001*** | 0.76 | < 0.001*** | 0.65 | < 0.001*** |
| Correlation with the Flow Index |  | r | -0.11 | -0.06 | 0.25 | -0.07 | 0.17 | -0.25 | -0.12 |
|  |  | p | 0.33 | 0.69 | 0.0030** | 0.55 | 0.11 | 0.0068** | 0.20 |
| Between the Difficulty Bins | Easy vs. Optimal | Z | -0.04 | -0.10 | -0.18 | 0.92 | 0.32 | 0.62 | -1.10 |
|  |  | p | 0.97 | 0.99 | 0.92 | 0.53 | 0.75 | 0.67 | 0.32 |
|  | Optimal vs. Hard | Z | 0.41 | -0.78 | 1.26 | -1.39 | 0.39 | -1.00 | 2.37 |
|  |  | p | 0.79 | 0.60 | 0.31 | 0.35 | 0.75 | 0.68 | 0.054 |
|  | Easy vs. Hard | Z | 0.69 | -1.90 | 1.58 | -0.82 | 0.40 | -0.72 | 1.13 |
|  |  | p | 0.67 | 0.096 | 0.25 | 0.56 | 0.79 | 0.64 | 0.32 |
| Success vs. Failure |  | Z | 1.74 | -2.31 | 0.92 | 1.06 | 0.72 | -1.28 | -1.69 |
|  |  | p | 0.18 | 0.021* | 0.45 | 0.48 | 0.65 | 0.60 | 0.17 |

#### 3.1.2. Associations of FMT activity with psychological and behavioral indices

To confirm consistency with previous studies reporting the relationships of FMT activity with psychological and behavioral measurements, the present study analyzed how the FMT ratio changed with the flow index, difficulty index, and success of gameplay.

First, given that FMT activity is a measure for estimating the flow state (Ewing, Fairclough, and Gilleade 2016; Katahira et al. 2018; Knierim et al. 2018; Farrugia et al. 2021), it should positively correlate with the flow index. However, the positive correlation between the FMT ratio and the flow index was significant in only one of our seven participants (ID2; game title: Smash Bros.) (Figure 2, Table 3), and another participant showed a significant negative correlation (ID6; game title: VALORANT) (Figure 2, Table 3). This result contradicts the conventional assumption that greater FMT activity occurs when a deeper flow state is experienced. Second, previous research has indicated that FMT activity during task implementation is greater when the difficulty of the task is optimal for a given individual compared with when it is either too easy or too hard (Ewing, Fairclough, and Gilleade 2016; Katahira et al. 2018; Knierim et al. 2018; Farrugia et al. 2021). In contrast with these previous findings, none of our participants showed a significantly higher FMT ratio in the trials of optimal difficulty (Figure 3A, Table 3). Third, to confirm whether enhanced top-down cognitive function induced high performance in games, the relationship between the FMT ratio and performance was investigated. However, no participant showed a significantly larger FMT ratio in the success bin, and one participant showed a significantly smaller FMT ratio in that bin (Figure 3B, Table 3). It is thus suggested that FMT dynamics do not correspond to the outcomes of games.

**Figure 2.**
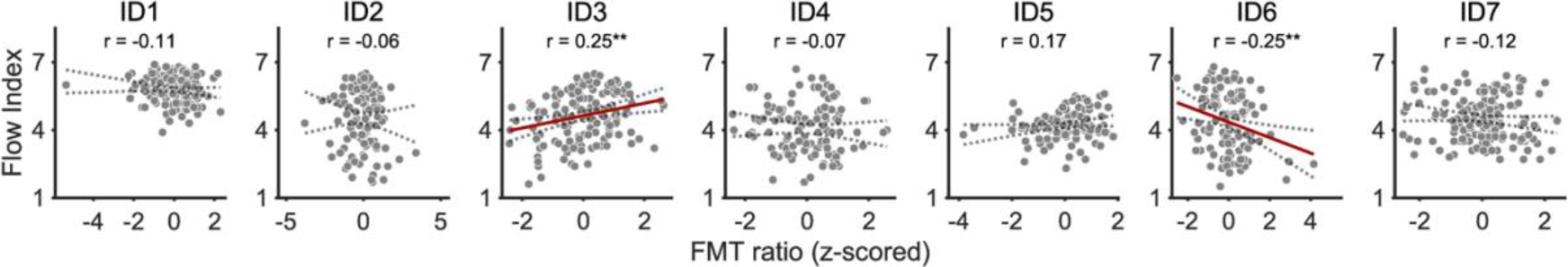
Correlation between the frontal-midline theta (FMT) ratio (z-scored) and flow scores. Each dot denotes one trial. Red lines represent regression lines. Broken lines denote the confidence intervals of the r-values. The false discovery rate was corrected for multiple comparisons. **p < 0.01.

**Figure 3.**
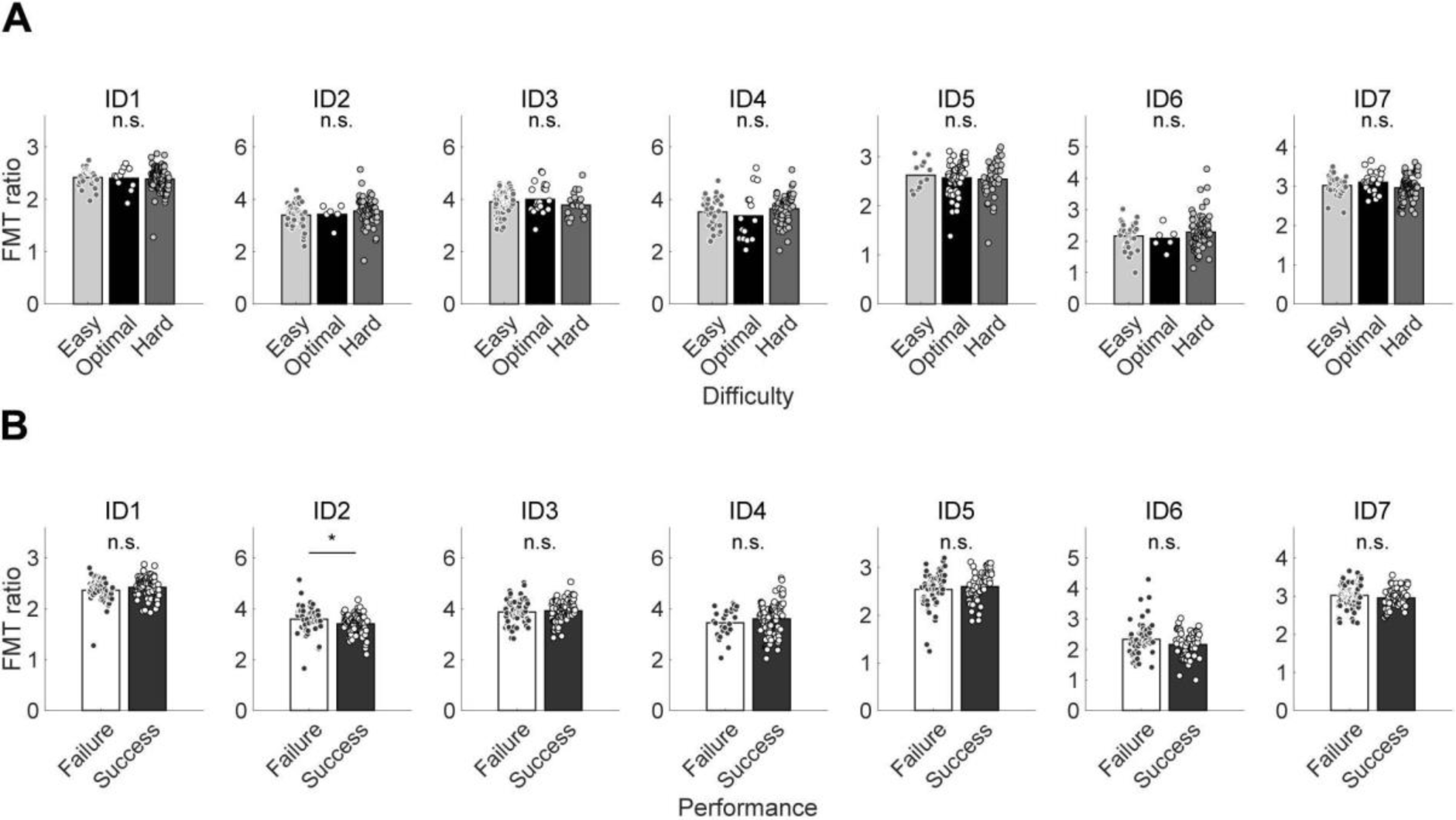
The frontal-midline theta (FMT) ratio according to subjective difficulty **(A)** and game performance **(B)**. Bars represent the FMT ratio for the easy, optimal, and hard bins in **(A)** and failure and success bins in **(B)**. Each dot denotes one trial. The false discovery rate was corrected for multiple comparisons. *p < 0.05; n.s., not significant.

### 3.2. Associations between flow states and other indices

The present study also examined the associations between flow states and other indices. The flow index was significantly correlated with RRI_min_ in four of the seven participants (Figure 4, Table 4). In other words, these four participants showed faster heart rates in trials with a deeper flow experience, consistent with previous studies (de Manzano et al. 2010; Gaggioli et al. 2013). As for the relationship between flow experience and game performance, the flow index was significantly higher in the successful trials compared with the failed ones in all seven participants (Supplementary Figure S1, Table S1). By contrast, RRI_min_ did not differ between the successful and failed trials in six out of the seven participants (Supplementary Figure S2, Table S1). Regarding the interaction between flow experience and subjective difficulty, the flow index was significantly lower for the hard bins than the other bins in four of the seven participants, while it was highest for the easy bin in one participant and did not differ among bins in the other two participants (Supplementary Figure S3, Table S1). Meanwhile, RRI_min_ was highest for the easy bins in three of the seven participants, while it was lowest for the optimal bin in one participant and did not differ among bins for the other three participants (Supplementary Figure S4, Table S1).

**Figure 4.**
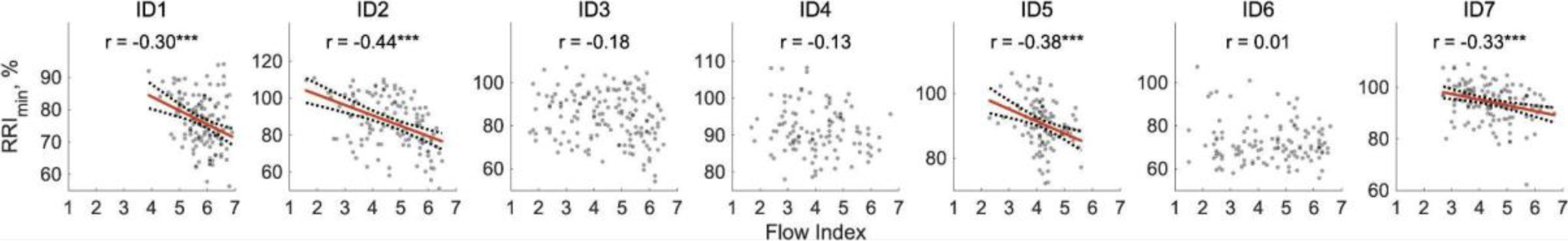
Correlation between flow index and RRI_min_. Each dot denotes one trial. Red lines represent regression lines. Broken lines denote the confidence intervals of the r-values. The false discovery rate was corrected for multiple comparisons. ***p < 0.001.

**Table 4.** Results of the correlation analysis between flow index and RRI_min_. The false discovery rate was corrected for multiple comparisons. ***p < 0.001.

|  | ID1 | ID2 | ID3 | ID4 | ID5 | ID6 | ID7 |
| --- | --- | --- | --- | --- | --- | --- | --- |
| $r$ | -0.30 | -0.44 | -0.18 | -0.13 | -0.38 | 0.01 | -0.33 |
| $p$ | < 0.001*** | < 0.001*** | 0.067 | 0.31 | < 0.001*** | 0.89 | < 0.001*** |

## 4. Discussion

The present study examined the relationship between the subjective experience of the flow state and FMT activity, which has been considered a hallmark of the flow state in previous research (Ewing, Fairclough, and Gilleade 2016; Katahira et al. 2018; Knierim et al. 2018; Farrugia et al. 2021). Participants played their favorite video games (in which they were also most proficient) at the optimal challenge level. EEG, ECG, and psychological questionnaire responses were recorded during more than 100 gameplay sessions per participant. Similar to previous studies, five out of seven participants showed higher FMT power when playing games compared with the baseline phase. Surprisingly, however, in most participants (six out of seven), there was no positive correlation between FMT power while playing games and the degree of subjective flow experience, inconsistent with previous findings.

### 4.1. Difference in FMT activity between the gameplaying and resting states

The increases in FMT power seen during gameplay in this study suggest activation of the frontal lobe. This is consistent with previous findings that FMT activity increases during cognitive tasks (Onton, Delorme, and Makeig 2005; Popov et al. 2018). However, in two of our participants, FMT activity did not increase during gameplay, indicating individual differences in the increase in FMT activity during cognitive tasks (Jensen et al. 2002).

### 4.2. Correlation of FMT activity with flow scores and performance

The present study employed an experiment that addressed the four major methodological issues highlighted in previous studies (Alameda, Sanabria, and Ciria 2022; Grice et al. 2020). First, instead of unfamiliar, non-motivational tasks, we used participants’ favorite games (in which they were also most proficient) to induce motivation and facilitate the experience of a flow state, as well as to allow for measurement of flow and neural activity in a more competitive situation. Second, previous studies have directly related the neural activity obtained with different task difficulty levels to the flow state, and they could not exclude the possibility that this neural activity was generated by differences in task difficulty. However, this task difficulty manipulation ignored the fluctuation of flow states when task difficulty was unchanged (e.g., during swimming, darts, and video games with opponents matched for ability). Thus, in the present study, we employed a task design that allowed us to directly relate the variability in subjective ratings of flow states to neural activity by keeping the task difficulty as constant as possible. Nevertheless, there was intra-individual variability in the FSS scores collected after each match. This is reasonable because the flow state is considered to be influenced not only by external factors, such as task difficulty, but also by internal factors fluctuating within and across trials, such as the individual’s motivation (Moneta and Csikszentmihalyi 1999; Gold and Ciorciari 2020). In addition, the negative correlation between RRI_min_ and the flow index in four out of seven participants suggests that parasympathetic activity can vary when the level of difficulty is constant. Third, the duration of the experiments in previous studies was sufficient to observe the relatively rare phenomenon of flow states. In this study, more than 100 game play sessions were completed over 10 days by each participant to increase the likelihood of capturing high flow states. Finally, in contrast to most previous studies, which analyzed flow states at the group level, the present study adopted a within-individual statistical approach. Because psychological states and emotions exhibit marked individual differences, a method that averages data across individuals would produce misleading resulting by ignoring the existence of individual differences (Fischer, Nilsson, and Ebner 2023; Smith and Little 2018). In this study, statistical analyses were conducted for each individual, and trends across individuals were then explored. This allowed us to more accurately capture the true nature of flow states, which vary widely among individuals. By adopting such an experimental paradigm, we could examine the relationship between the flow state and FMT activity, resulting in findings differing from those of previous research.

The aim of this study was to investigate whether FMT activity is the neural correlate of the flow state. FMT activity reflects top-down cognitive control by the frontal lobes (Cavanagh and Frank 2014; Berger et al. 2019; Pavlov and Kotchoubey 2020) and has been considered a hallmark of the flow state (Ewing, Fairclough, and Gilleade 2016; Katahira et al. 2018; Knierim et al. 2018; Farrugia et al. 2021). In the present study, however, most (six of seven) participants did not show significant positive correlations between FMT activity and subjective flow intensity. This is a surprising result contradicting the assumption that top-down cognitive control, reflected in FMT activity, is the neural mechanism underlying the flow state. The present study strongly suggests that FMT activity is not the primary neural basis for the flow state.

If top-down cognitive control is not involved in the flow state, what neural mechanisms might be involved? Some studies have proposed that downregulation of the default mode network (DMN) is related to the flow state (Alameda, Sanabria, and Ciria 2022; van der Linden, Tops, and Bakker 2021). The DMN is a complex whole-brain network involved in self-referential processing and autobiographical memory (Davey, Pujol, and Harrison 2016; Smallwood et al. 2021). Given these characteristics of the DMN, inhibition of DMN activity might lead to fewer worries or self-focused thoughts during the flow state. To clarify whether DMN activity correlates with the flow state, further study is needed.

### 4.3. Relationships of FMT with subjective difficulty and performance

We compared FMT ratios among trials classified in terms of difficulty as “optimal” (score of 5 on the difficulty index), “too easy,” or “too hard.” In contrast to previous studies, we did not observe a significant increase in the FMT ratio under the optimal difficulty condition.

This result is probably attributable to our unique experimental design controlling task difficulty across trials. In this study, an online matching system was implemented so that each participant’s skill corresponded to that of their opponent. Therefore, the difficulty index could reflect smaller differences in difficulty compared with previous studies, which likely explains the inconsistency between the present study and previous ones in terms of the relationship between subjective difficulty and FMT activity.

### 4.4. Limitations

The present study employed an online matching system to control the difficulty level across trials, avoided the possibility of task adaptation, and required participants to try their best to succeed in as many trials as possible. However, because both the participants and their opponents did not always play their best in every match, variation in task difficulty was inevitable. Nevertheless, considering the absence of significant differences in the FMT ratio among the difficulty bins (Figure 4B), the variation in difficulty across trials was sufficiently small to not modulate frontal lobe activities.

In the present study, the participants played their favorite games, which were also those with which they felt most confident; thus, there were differences among participants in the genres of games played. We observed enhanced FMT activity in the occipital region during gameplay compared with the baseline phase in two of the seven participants (ID5 and ID6). Considering that both of these participants played shooter games, the neural correlates for the flow state may differ among game genres. This supposition pertains mainly to the present study, where the neural activity associated with different games was evaluated using an identical procedure, but it may also explain the inconsistencies among previous findings et al. 2018; Farrugia et al. 2021). It is suggested that the neural basis for the flow state, i.e., the representative brain regions or frequency bands involved in that state, might be task-specific and variable among individuals.

## 5. Conclusion

The present study examined whether the subjective flow state correlates with FMT activity. We employed a task design that solved the methodological problems in previous studies investigating the neural correlates of the flow state, and we performed measurements over 100 gameplay sessions for each of seven participants. The results showed no positive correlation between FMT activity and subjective flow level in six out of seven participants, contrary to previous reports. Our results challenge the conventional view that frontal lobe activity, as reflected in FMT activity, is instrumental in achieving the flow state.

## 6. Data availability

The dataset of EEG, ECG, and FSS and custom-build MATLAB scripts for the present study was available from https://zenodo.org/doi/10.5281/zenodo.12662201.

## 7. Author contributions

Conceptualization, H.S. and T.I.; Methodology, T.I. and R.Y.; Investigation, H.S., T.I., and R.Y.; Data acquisition, H.S., T.I., and R.Y.; Data analyses, T.I. and R.Y.; Statistical analyses, R.Y.; Visualization, T.I. and R.Y.; Writing – original draft, H.S., T.I., and R.Y.; Writing – review & editing, H.S., T.I., R.Y., and J.U.; Supervision, J.U.; Funding acquisition, H.S. and J.U. All authors contributed to the article and approved the submitted version.

## Acknowledgments

This work was supported by a designated from Living Platform, Ltd., Japan to JU and by funding from HAYAO NAKAYAMA Foundation for Science & Technology and Culture, provided to HS. We thank Ms. Tomomi Hamaoka for her secretarial assistance. We also thank Ms. Mone Fujita for her contribution to the data analysis, and Mr. Aoi Seki and all other members of our laboratory for their insightful comments on our work. Finally, we thank Michael Irvine, PhD, from Edanz (https://jp.edanz.com/ac) for editing a draft of this manuscript.

