## Appendices for "The flow state is not accompanied by frontal-midline theta activity: An EEG investigation of more than 700 video gameplay sessions"

### Flow short scale

The FSS questionnaire used in Engeser and Rheinberg (2008) was transcribed into Japanese. The 10 items evaluating the subjective flow experience are as follows: (1) I feel just the right amount of challenge; (2) My thoughts/activities run fluidly and smoothly; (3) I don't notice time passing; (4) I have no difficulty concentrating; (5) My mind is completely clear; (6) I am totally absorbed in what I am doing; (7) The right thoughts/movements occur of their own accord; (8) I know what I have to do each step of the way; (9) I feel that I have everything under control; and (10) I am completely lost in thought. The three items evaluating the subjective difficulty and challenge–skill balance are as follows: (1) Compared to all other activities which I partake in, this one is...; (2) I think that my competence in this area is...; and (3) For me personally, the current demands are...

### Supplementary materials

Table S1. Flow index and ratio of the minimal R–R interval in each trial to that of the corresponding baseline ($RRI_{min}$) according to difficulty and performance bin. The false discovery rate was corrected for multiple comparisons. *p < 0.05; **p < 0.01; ***p < 0.001.

| Bins | | ID1 | ID2 | ID3 | ID4 | ID5 | ID6 | ID7 |
|---|---|---|---|---|---|---|---|---|
| The Flow Index | Z | -5.39 | -6.26 | -7.63 | -5.44 | -5.19 | -5.36 | -6.14 |
| Success vs. Failure | p | < 0.001*** | < 0.001*** | < 0.001*** | < 0.001*** | < 0.001*** | < 0.001*** | < 0.001*** |
| $RRI_{min}$ | Z | 0.69 | 0.04 | -1.37 | 1.90 | 1.13 | -0.76 | 4.20 |
| Success vs. Failure | p | 0.61 | 0.97 | 0.28 | 0.17 | 0.43 | 0.67 | < 0.001*** |
| The Flow Index | Z | 0.92 | -0.81 | 1.40 | 0.34 | 0.83 | 0.25 | 1.82 |
| Easy vs. Optimal | p | 0.54 | 0.62 | 0.30 | 0.78 | 0.61 | 0.86 | 0.15 |
| The Flow Index | Z | 0.36 | 1.96 | -0.16 | 2.97 | 4.59 | 1.68 | -0.63 |
| Optimal vs. Hard | p | 0.77 | 0.093 | 0.87 | 0.0030** | < 0.001*** | 0.35 | 0.53 |
| The Flow Index | Z | 2.03 | 2.56 | 3.73 | 4.05 | 3.51 | 2.73 | 1.53 |
| Easy vs. Hard | p | 0.11 | 0.010* | < 0.001*** | < 0.001*** | < 0.001*** | 0.0064** | 0.19 |
| $RRI_{min}$ | Z | 2.87 | 2.35 | 0.95 | 1.77 | 2.58 | 0.93 | -0.67 |
| Easy vs. Optimal | p | 0.0041** | 0.019* | 0.47 | 0.19 | 0.0099** | 0.59 | 0.54 |
| $RRI_{min}$ | Z | -1.16 | -0.28 | 0.22 | -2.28 | -3.18 | -0.36 | 1.92 |
| Optimal vs. Hard | p | 0.41 | 0.90 | 0.95 | 0.086 | 0.0015** | 0.83 | 0.14 |
| $RRI_{min}$ | Z | 3.58 | 2.42 | 2.43 | -0.65 | 0.41 | 1.21 | 1.21 |
| Easy vs. Hard | p | < 0.001*** | 0.016* | 0.015* | 0.60 | 0.85 | 0.56 | 0.31 |

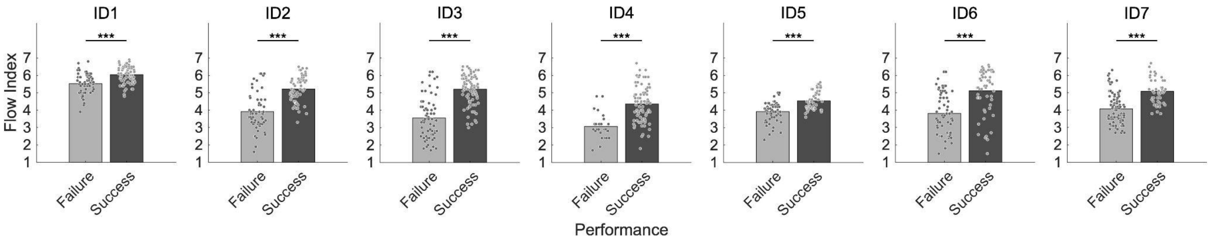


Figure S1. Flow index according to performance. Light and dark bars represent the flow index for the failure and success bins, respectively. Each dot denotes one trial. The false discovery rate was corrected for multiple comparisons. ***p < 0.001; n.s., not significant.

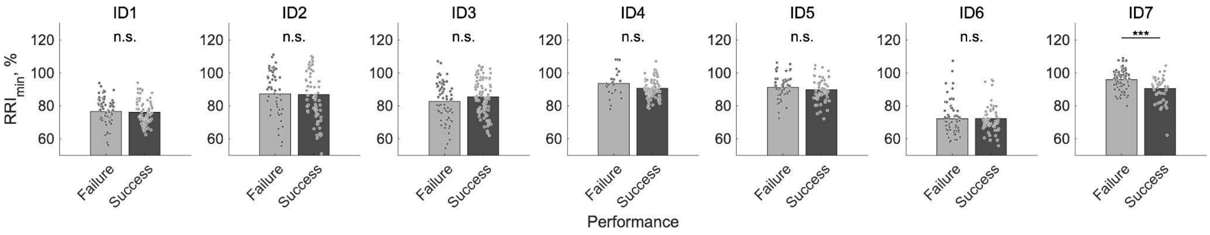


Figure S2. Ratio of the minimal R–R interval in each trial to that of the corresponding baseline ($RRI_{min}$) according to performance. Light and dark bars represent the $RRI_{min}$ for the failure and success bins, respectively. Each dot denotes one trial. The false discovery rate was corrected for multiple comparisons. ***p < 0.001; n.s., not significant.

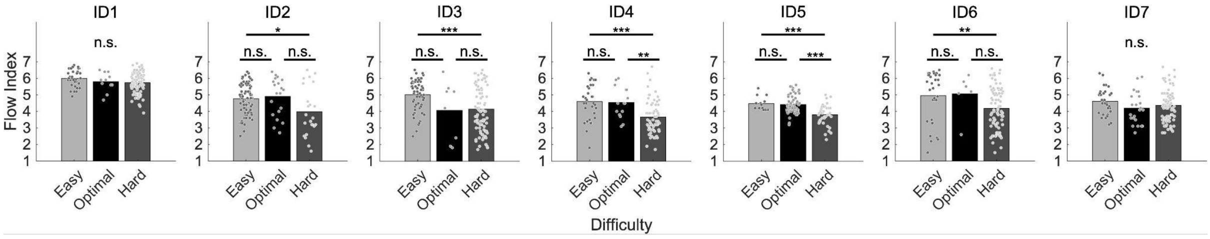


Figure S3. Flow index according to subjective difficulty. Bars represent the flow index for the easy, optimal, and hard bins. Each dot denotes one trial. The false discovery rate was corrected for multiple comparisons. *p < 0.05; **p < 0.01; ***p < 0.001; n.s., not significant.

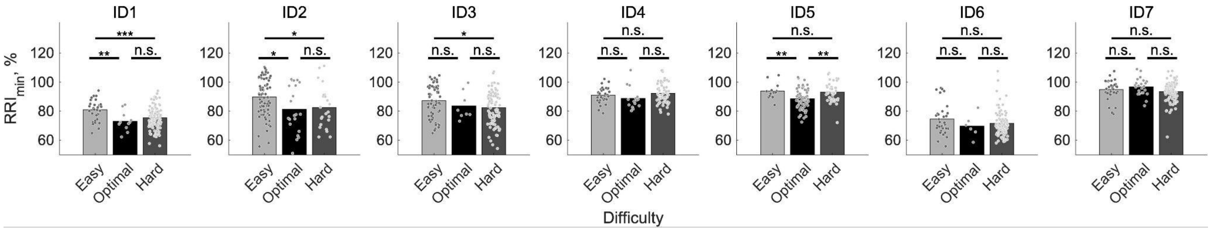


Figure S4. Ratio of the minimal R–R interval in each trial to that of the corresponding baseline ($RRI_{min}$) according to subjective difficulty. Bars represent the $RRI_{min}$ for the easy, optimal, and hard bins. Each dot denotes one trial. The false discovery rate was corrected for multiple comparisons. *$p < 0.05$; **$p < 0.01$; ***$p < 0.001$; n.s., not significant.
